# Quantitative assessment of constitutive G protein-coupled receptor activity with BRET-based G protein biosensors

**DOI:** 10.1101/2021.02.05.429900

**Authors:** Hannes Schihada, Rawan Shekhani, Gunnar Schulte

## Abstract

Heterotrimeric G proteins constitute the primary transducers of G protein-coupled receptor (GPCR) signaling. Besides mediating ligand-induced GPCR activation, G proteins transduce basal levels of activity in various physiological and pathophysiological settings evoked by constitutively active, native GPCRs or disease-related receptor mutants. Several generations of optical biosensors were developed and optimized to monitor GPCR ligand-induced G protein activation, however, quantitative approaches to detect constitutively active GPCRs are not available. Here, we designed and validated a set of eight bioluminescence-resonance-energy-transfer (BRET)-based G protein sensors, covering all four major families of G proteins, and established a protocol to identify constitutive GPCR/G protein signaling in living cells. These sensors rely on the encoding of all three G protein subunits on a single plasmid, enabling their cellular expression at desired relative levels and resulting in reduced signal variability in mammalian cells. Based on this sensor platform, we further present here an experimental protocol to quantify constitutive signaling of native and mutated GPCRs through these heterotrimeric transducers. This approach will aid in the characterization of constitutively active GPCRs and the exploration of their role in health and disease.

**One Sentence Summary:** This Resource article describes the validation of a biophysical approach to directly assess the constitutive signaling activity of G protein-coupled receptors through heterotrimeric G proteins in living cells using optical biosensors.

## Introduction

Heterotrimeric G proteins represent the prime intracellular transducer proteins of G protein-coupled receptors (GPCRs), which constitute the largest family of membrane targets for approved drugs (*1*). G proteins are plasma membrane-anchored complexes composed of three distinct subunits, named in order of their respective molar masses G_α_, G_β_ and G_γ_, and grouped into four major families, G_s_, G_i/o_, G_q/11_ and G_12/13_, based on the signaling pathways promoted by the G_α_ subunit (*2*). These three subunits form inactive heterotrimeric holoenzymes with guanosine diphosphate (GDP) bound to G_α_. Ligand-induced GPCR activation and subsequent GPCR/G protein interaction catalyzes GDP exchange for guanosine triphosphate (GTP) and complex dissociation into active G_α_ and G_βγ_ molecules, further relaying the signal to a plethora of cellular effectors (*2–4*). However, the concept of G_α_-G_βγ_ dissociation as a consequence of G protein activation has been challenged by previous studies (*5–8*) and remains a matter of debate (*9–11*).

In addition to agonist-induced receptor/G protein activation, numerous GPCRs exert tissue-specific, constitutive (i.e., agonist-independent) G protein activation. Moreover, excessive signal transduction through gain-of-function mutations in GPCR genes is implicated in various human disorders (*12–16*). Since the late 80s, investigating constitutive GPCR activity has mostly relied on either (i) laborious biochemical quantification of GTPase activity or (ii) measurements of downstream effector activity (e.g. adenylyl cyclases (AC) activated and inhibited by G_s_ and G_i/o_, respectively), intracellular second-messenger levels (e.g. cyclic adenosine monophosphate, cAMP, downstream of G_αs_ and G_αi/o_ and inositol phosphates, IP_1/2/3_, downstream of G_αq/11_ signaling) and other integrated cellular responses even further downstream of receptor/G protein interactions (*17–27*). However, assessment of constitutive activity of GPCR/G protein complexes based on intracellular second messenger levels is limited to subtypes of the G_s_ and G_q/11_ families, does not reveal functional selectivity within the same G protein family (e.g. G_q_ vs. G_15_), and often requires control experiments with inverse agonists, which are not available for many GPCRs. For these reasons, more proximal, direct and generalizable methods, applicable to all G protein families with subtype-specific resolution are required to accelerate the exploration of constitutive GPCR/G protein signaling and to allow for a comprehensive screening of basal GPCR activity for drug discovery.

Several generations of fluorescence- and bioluminescence-resonance-energy-transfer (FRET and BRET)-based G protein biosensors were developed since their initial description in the early 2000s (*5, 28–36*) and seminal work in identifying optimal labeling sites in individual G protein subunits, in particular in G_α_ (*8, 37*), has been done. The great majority of these sensors is based on tagging G_α_ with a FRET or BRET donor and either G_β_ or G_γ_ with an appropriate fluorescent protein serving as an energy acceptor. Building on these efforts in G protein biosensor developments, a more recent large-scale comparison of BRET donor (i) insertion sites within G_α_ and (ii) G_α_/G_β_/G_γ_ combinations resulted in a toolbox of sensitive G protein biosensors named TRUPATH (*37*). Despite the improved sensitivity of these biosensors in head-to-head comparisons with previous G protein BRET probes, measuring G protein activity still requires simultaneous host cell transfection with three plasmids, each encoding one of the synthetic G protein subunits. This limitation leads to inter-cell varying expression levels of the G protein subunits, which has a substantial impact on the assay’s sensitivity when submaximal effects, such as those arising from GPCR constitutive activity, are under investigation. Moreover, the necessity to co-transfect a mammalian cell population with three individual plasmids cannot ensure a balanced expression ratio of the three subunits, artificially changing the BRET signal of these intermolecular sensors and, ultimately, disguising the BRET responses induced by co-expression of a constitutively active GPCR.

For these reasons, we aimed to develop a novel set of BRET-based G protein activity sensors, that rely on the seminal developments of optical G protein probes, and overcome the limitations arising from multi plasmid co-transfection. This novel set of BRET sensors encoded on tricistronic plasmids exhibits improved sensitivity compared to their three-plasmid-based counterparts and allowed us to establish an experimental setup to assess the constitutive activity profiles of native receptors and disease-related GPCR mutants.

## Results

### Design of G protein activity sensors

To generate BRET-based biosensors that detect G protein activity upon GPCR-mediated G_α_-G_βγ_ dissociation (**Fig. 1A**), we (i) tagged G_α_ and G_γ_ with suitable energy transfer partners, and (ii) encoded these labeled constructs along with native G_β_ on a single plasmid (**Fig. 1B**). We took advantage of the seminal work in identifying optimal insertion sites in G_α_ proteins (*33, 37*) and tagged eight distinct G_α_ subunits, covering all four major G protein families, with the small and bright luciferase NanoLuciferase (Nluc) (*38*) at the recommended sites. G_γ_ subunits were N-terminally labeled with circularly permuted Venus (cpVenus^173^) since this fluorescent protein has served as a sensitive energy acceptor in previous FRET and BRET biosensors (*33, 39, 40*). The genes encoding these engineered G protein subunits were then combined on a single plasmid (**Fig. 1C**). G_β_ and G_γ_ cDNA were encoded as a single transcript, linking the subunits with an 18 amino acid viral T2A peptide sequence that is self-cleaved post-translationally in intact cells (*41*). Co-expression of Nluc-G_α_ was facilitated by an upstream internal ribosome entry site (IRES) (*41*), as employed previously for Gi1-3 and G13 FRET sensors (*31, 33*) in order to reduce expression level variability upon transfection into mammalian host cells.

**Fig. 1:**
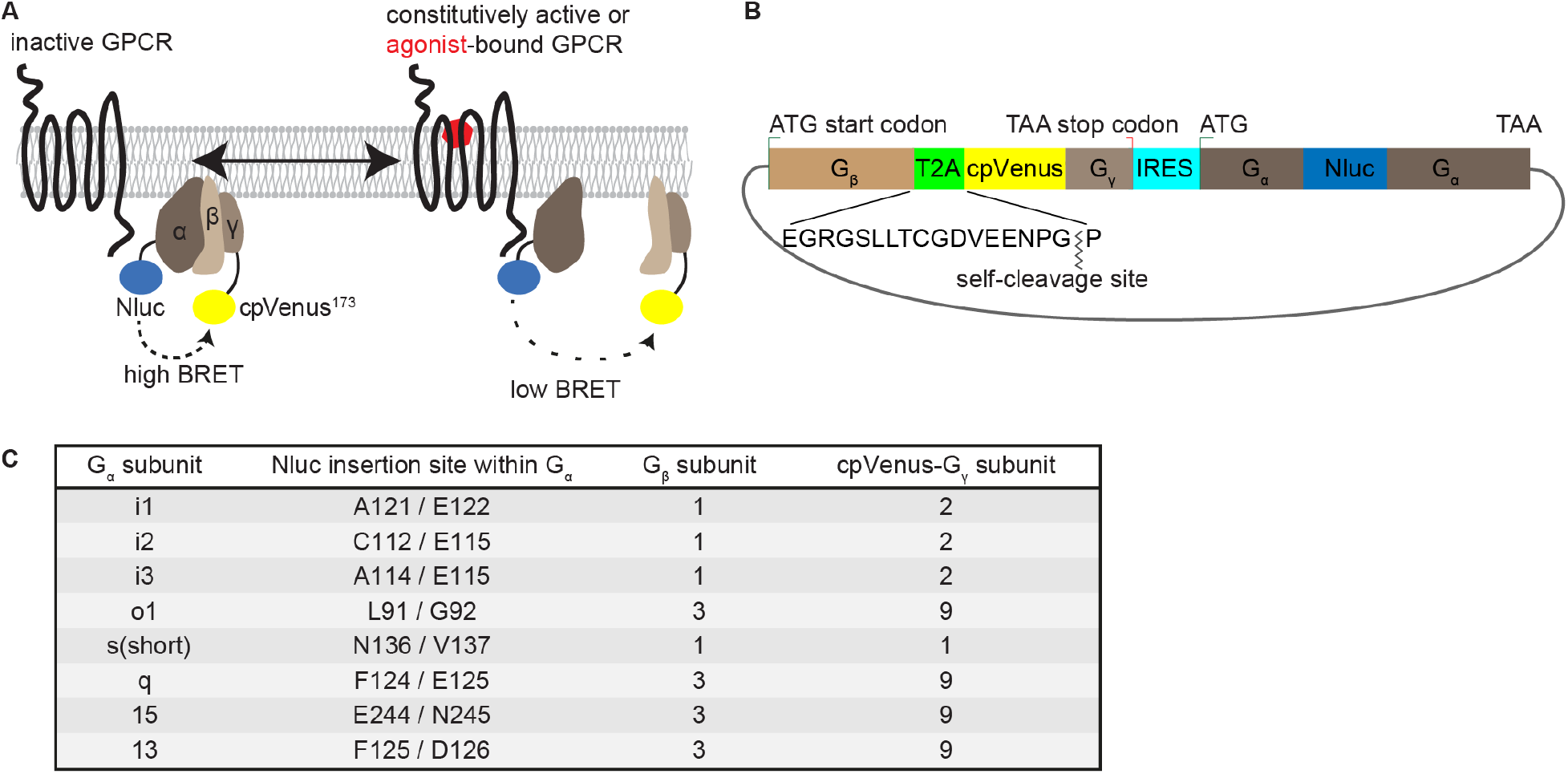
Design of BRET-based G protein activity sensors. **A)** Assay principle. **B**) Design of the tricistronic G protein sensor plasmid. **C**) Composition of tricistronic G protein sensor plasmids.

### Functional validation of G protein BRET sensors

First, we confirmed efficient co-expression of labeled G protein subunits upon transfection of these tricistronic plasmids in HEK293A cells by recording the luminescence emission spectra upon addition of the Nluc substrate furimazine (**Fig. S1**). Transfection with all eight biosensors resulted in luminescence spectra with a characteristic Nluc emission peak at ~ 450 nm and the cpVenus-related peak at ~ 530 nm, confirming (i) the successful co-expression of tagged G protein subunits upon transfection with a single plasmid and (ii) basal Nluc-to-cpVenus resonance energy transfer in each biosensor.

In order to verify the functionality of these biosensors to dynamically report G protein activity in living cells, we next co-transfected previously validated GPCRs along with the BRET sensors for their cognate G proteins (histamine H3 receptor (H3R) / G_i1_, G_i2_, G_i3_, G_o1_; β_2_-adrenergic receptor (β_2_AR) / G_s_; thromboxane A2 receptor (TBXA2R) / G_q_, G_15_, G_13_) (*42*). Agonist-induced activation of all eight receptor / G protein pairs resulted in significant time- and concentration-dependent BRET reductions, revealing activated receptor-mediated dissociation of Nluc-tagged G_α_ and cpVenus-tagged G_βγ_ subunits (**Fig. 2A-H**). The amplitude of the BRET response ranged from ~ −10% for G_s_ and G_15_ to ~ −35 % with G13, indicating distinct sensitivities of these biosensors and/or G_α_-specific underlying mechanisms following GPCR activation. The concentration-response curves of these BRET signals revealed ligand potencies that were in good agreement with values obtained with other receptor-specific readouts (**Fig. 2I-P**) (*28, 42, 43*), further confirming the reliability of this novel sensor platform to detect ligand- and GPCR-dependent signaling through individual effector proteins.

**Fig. 2:**
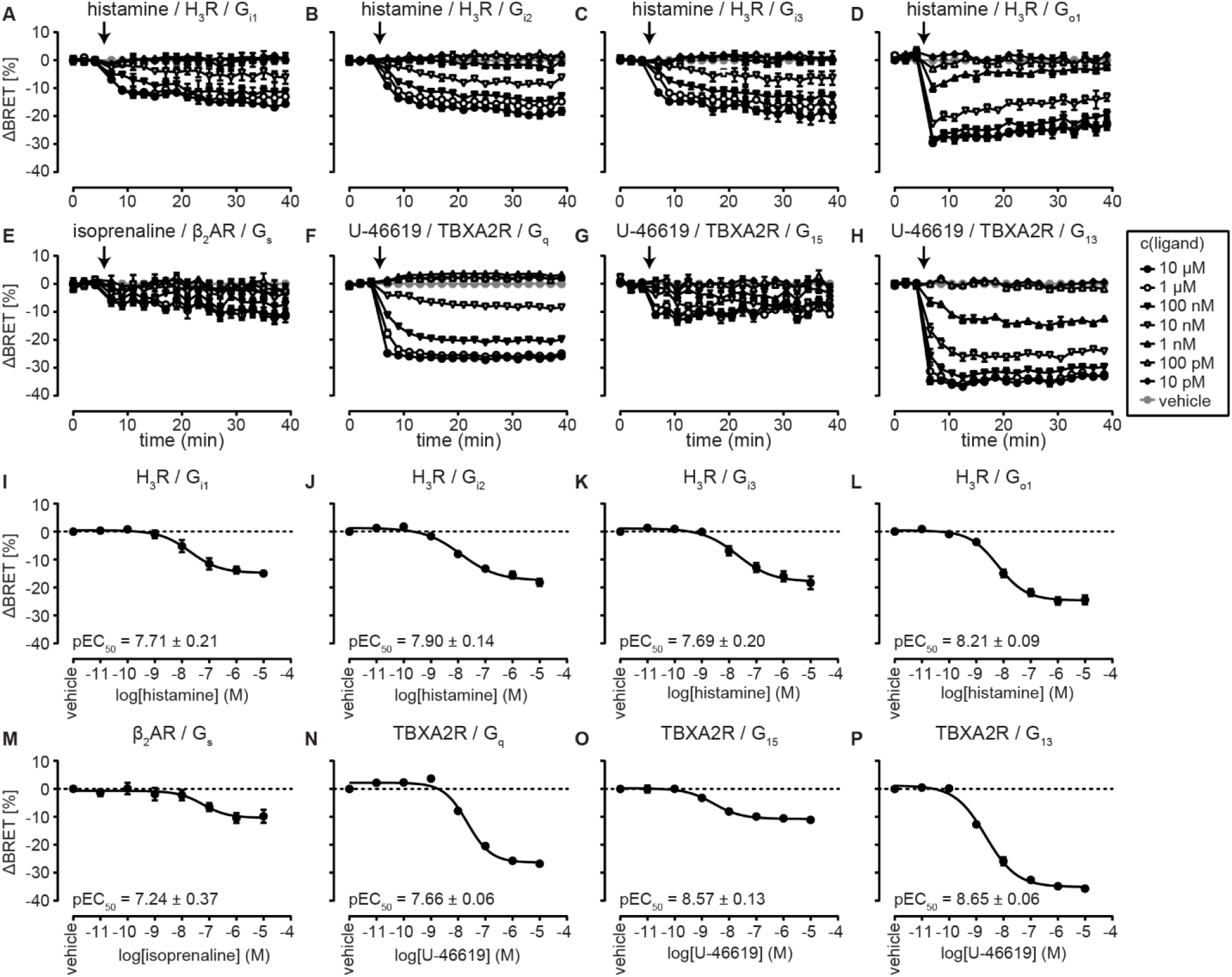
Functional validation of BRET-based G protein biosensors. **A-H)** ΔBRET time-courses of eight G protein BRET sensors upon GPCR agonist addition. **I-P**) Corresponding ΔBRET ligand concentration-response curves. All experiments were conducted in HEK293A cells transiently co-transfected with the indicated GPCR/G protein sensor pair and stimulated with the indicated GPCR ligand (arrow indicates time point for stimulation). Data show mean ± s.e.m. of three to four independent experiments.

### Superior sensitivity of the tricistronic sensor design

After confirming that these tagged G protein subunits are mutually expressed and report agonist-induced GPCR/G protein activation, we aimed to assess whether the tricistronic sensor design yields higher assay sensitivity compared to parallel transfection with three separate G protein subunits. We co-expressed the thromboxane A2 receptor (TBXA2R) along with either all three plasmids separately encoding the G13 sensor subunits (pcDNA-G_α13_-Nluc + pcDNA-G_β3_ + pcDNA-cpVenus-G_γ9_) (**Fig. 3A, Fig. S2A, B**), or with the tricistronic G13 sensor plasmid (**Fig. 3B, Fig. S2C, D**) and treated the cells with vehicle control or the TBXA2R agonist U-46619 (1μM) to calculate the resulting Z-factor (*44*). These experiments revealed a substantially higher average Z-factor and lower coefficient of variation (below 10%) with the tricistronic transfection system and demonstrate its improved, even high-throughput screening amenable, assay sensitivity, robustness and reproducibility (**Fig. 3C**).

**Fig. 3:**
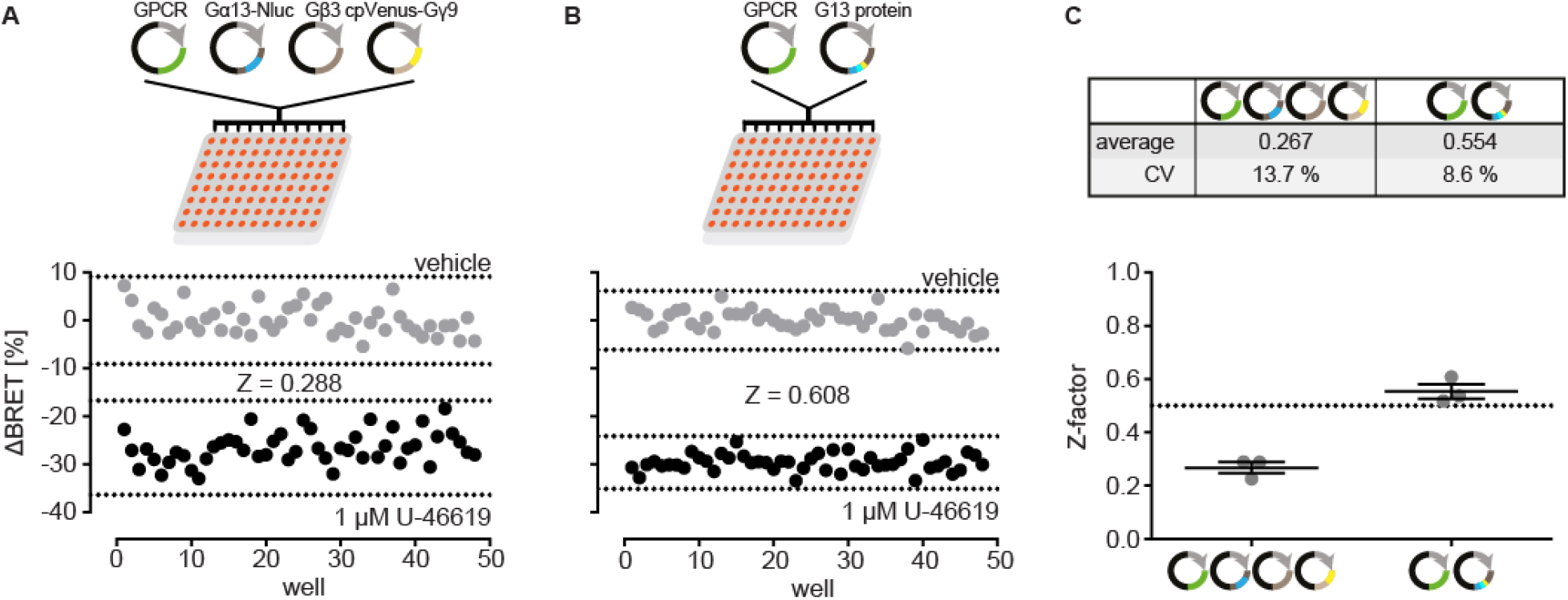
Improved assay sensitivity with tricistronic sensor design. **A)** Representative ΔBRET signals of the G_13_ BRET biosensor upon separate co-transfection with the three G protein subunits and TBXA2R. **B**) Representative ΔBRET signals of the G_13_ BRET biosensor upon transfection of the tricistronic G protein sensor plasmid along with TBXA2R. The dotted lines in (A) and (B) represent mean ± three-fold standard deviation of vehicle- and TBXA2R agonist (1 μM U46619)-induced BRET changes. **C**) Mean ± s.e.m. Z-factors of three independent G_13_ sensor experiments upon transfection with the separate (left) or tricistronic (right) sensor plasmids and TBXA2R. All experiments were conducted in transiently transfected HEK293A cells.

### Luminescence/BRET correlation

The superior sensitivity of these novel G protein biosensors tempted us to explore whether these optical tools are able to detect dissociation of heterotrimeric G proteins upon co-expression of a constitutively active GPCR, i.e. without the addition of a cognate GPCR agonist.

In theory, co-expression of a constitutively active receptor would result in a reduced G protein BRET ratio compared to empty vector-transfected control cells, independently from the G protein sensor’s expression level (model in **Fig. 4A**). However, both technical and biological factors can contribute to increasing or decreasing BRET ratios over varying absolute G protein sensor emission intensities within the same biological sample (model in **Fig. 4B**). The former include emission wavelength-dependent sensitivities of photon-collecting, -counting and - converting (into electrical signals) optical detectors in commercially available luminescence plate readers (*45*). For instance, a higher sensitivity in detecting increases in emission intensity in the cpVenus channel would artificially elevate the sensor’s BRET ratio over increasing relative luminescence units (e.g. resulting from varying sensor expression levels or cell densities). On the other hand, biological factors affecting G protein sensor activity (and therefore its BRET ratio) include relative expression levels of the overexpressed G proteins to endogenously expressed interaction partners (such as constitutively active GPCRs, native G protein subunits associating with exogenous labeled sensor molecules, non-GPCR activators of G protein signaling or GTPase-activating proteins, e.g. regulators of G protein signaling, RGS; (*46, 47*)). Furthermore, a high density of overexpressed G protein sensors can result in BRET from one Nluc molecule to multiple BRET acceptors and the relative G protein sensor / GPCR expression levels will presumably affect the calculated BRET ratio (i.e. the higher the G protein excess, the smaller the BRET reduction due to const. GPCR activity) (model in **Fig. 4C**). These underlying confounders would hamper a direct comparison of G protein BRET ratios in GPCR- vs. control-transfected samples for the assessment of constitutive GPCR activity (see blue encircled data points in **Fig. 4B, C**).

**Fig. 4:**
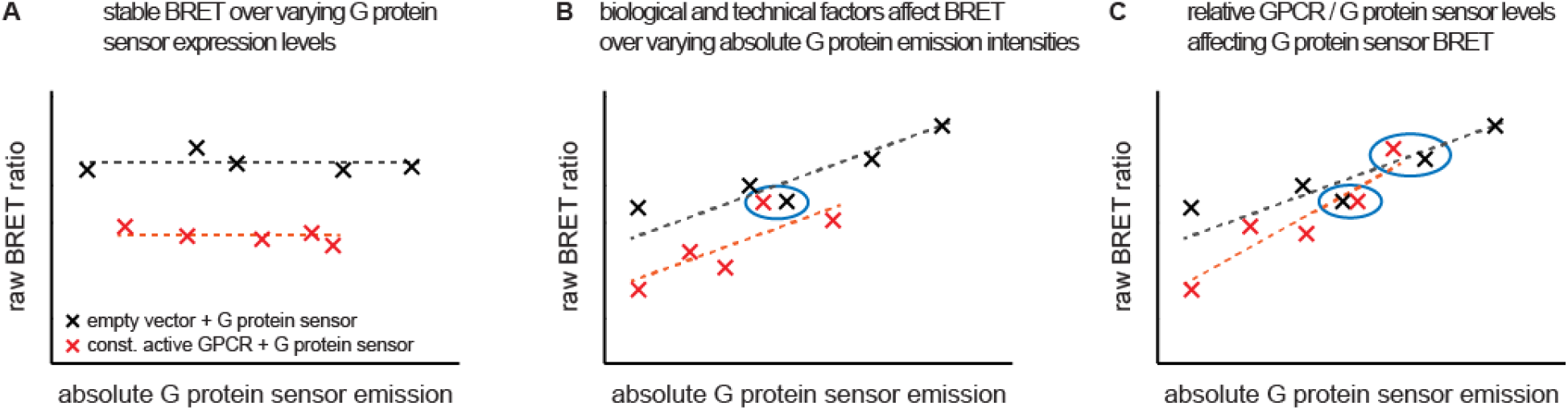
Theoretical models for correlation of the measured BRET ratio and G protein sensor emission. **A)** Scheme for luminescence intensity-independent BRET behavior (slopes equal to 0). Co-transfection of constitutively active GPCRs results in a reduced BRET ratio over the entire range of G protein sensor intensities. **B**) Direct linear correlation of G protein sensor emission and the measured BRET ratios due to technical and biological factors affecting photon detection and conversion, and modulating G protein activity, respectively. Expression of constitutively active GPCRs results in a parallel linear correlation with reduced BRET ratios at similar G protein sensor emission intensities. **C**) G protein sensor co-transfection with control vs. constitutively active GPCRs results in linear correlation with distinct slopes. The molecular GPCR:G protein sensor ratio, which is highest at low G protein sensor intensities, elicits a steeper linear fit compared to the negative control model.

To overcome these obstacles, we set out to first evaluate experimentally how BRET changes over varying G protein sensor luminescence intensities (**Fig. 5A-H**). As expected, BRET was not stable over the tested range of emission intensities and resulted in a direct linear correlation between relative Nluc light units (representing the absolute G protein sensor levels per well) and the corresponding BRET ratio (assessed through runs test for linearity, **Table S1**). Of note, technical settings of the luminescence measurement (e.g. channel bandwidth and photon number multiplicators “gain”) determine the parameters of this linear BRET/Nluc correlation.

**Fig. 5:**
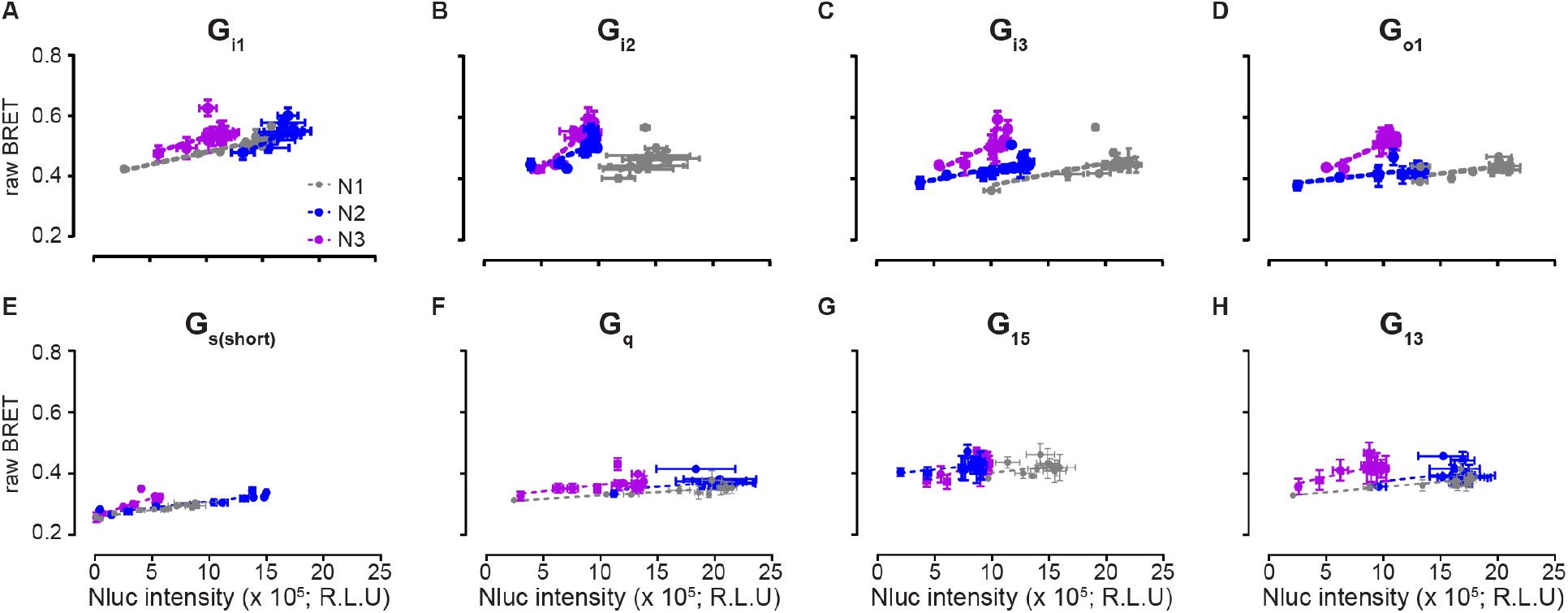
Experimental validation of the correlation between measured BRET ratios and G protein sensor emission. BRET over Nluc luminescence plots of the indicated G protein sensors. All experiments were conducted in HEK293A cells transiently transfected with increasing amounts of the tricistronic G protein sensors. Data show mean ± s.d. of three individual experiments and fitted to a linear correlation. Runs test was conducted to confirm linear correlation between BRET and Nluc emission intensity (see Table S1).

### Assessment of constitutive GPCR signaling through heterotrimeric G proteins

In order to evaluate the fidelity of the G protein biosensors to detect constitutively active GPCRs, we co-transfected four previously confirmed, constitutively active GPCR/G protein pairs in HEK293A cells: H_3_R with G_i1_ (*48–52*), the Cannabinoid receptor 1 (CB_1_) with G_i3_ (*53–56*), β_2_AR with G_s_ (*57–60*), and TBXA2R with G_q_ (*61–63*). By plotting the BRET ratios of co-transfected samples as a function of the sensors’ relative light units (in the Nluc emission channel), we determined the Y-intercept of each fit. This extrapolated BRET value, henceforth referred to as ‘BRET_0_’, allows us to correct for varying luminescence intensities and to compare G protein activity at a fixed (virtual) sensor expression level (**Fig. 6A-D**). This biophysical approach confirmed constitutive signaling of all four GPCRs through their cognate G proteins (**Fig. 6E – H**), underlining the reliability of our method to detect constitutively active GPCRs. In contrast, comparing the BRET ratios of receptor- vs. pcDNA-transfected samples without correction for distinct sensor emission intensities would not reveal the constitutive signaling activity of H_3_R and β_2_AR (**Fig. S3**). In addition to these findings with known constitutively active GPCRs, experiments with the G_i/o_-coupled α_2A_-adrenergic receptor (α_2A_AR) exhibited a significantly reduced BRET_0_ value only with G_o1_ but not with any other G protein biosensor (**Fig. 6I, Fig. S4**). These data resemble the findings of a previous study where α_2A_AR signaled constitutively through co-transfected G_αo_ (*64*), and highlight the specificity of this method in detecting constitutively active GPCRs with a low number of false-positive results. To further confirm our results of constitutive activity GPCR/G protein activity, we applied the CB_1_ receptor inverse agonist rimonabant to CB_1_ receptor/G_i3_-transfected cells and recorded a rapid, concentration-dependent increase in BRET, confirming the re-assembly of a pre-activated heterotrimer upon GPCR de-activation (**Fig. S5**).

**Fig. 6:**
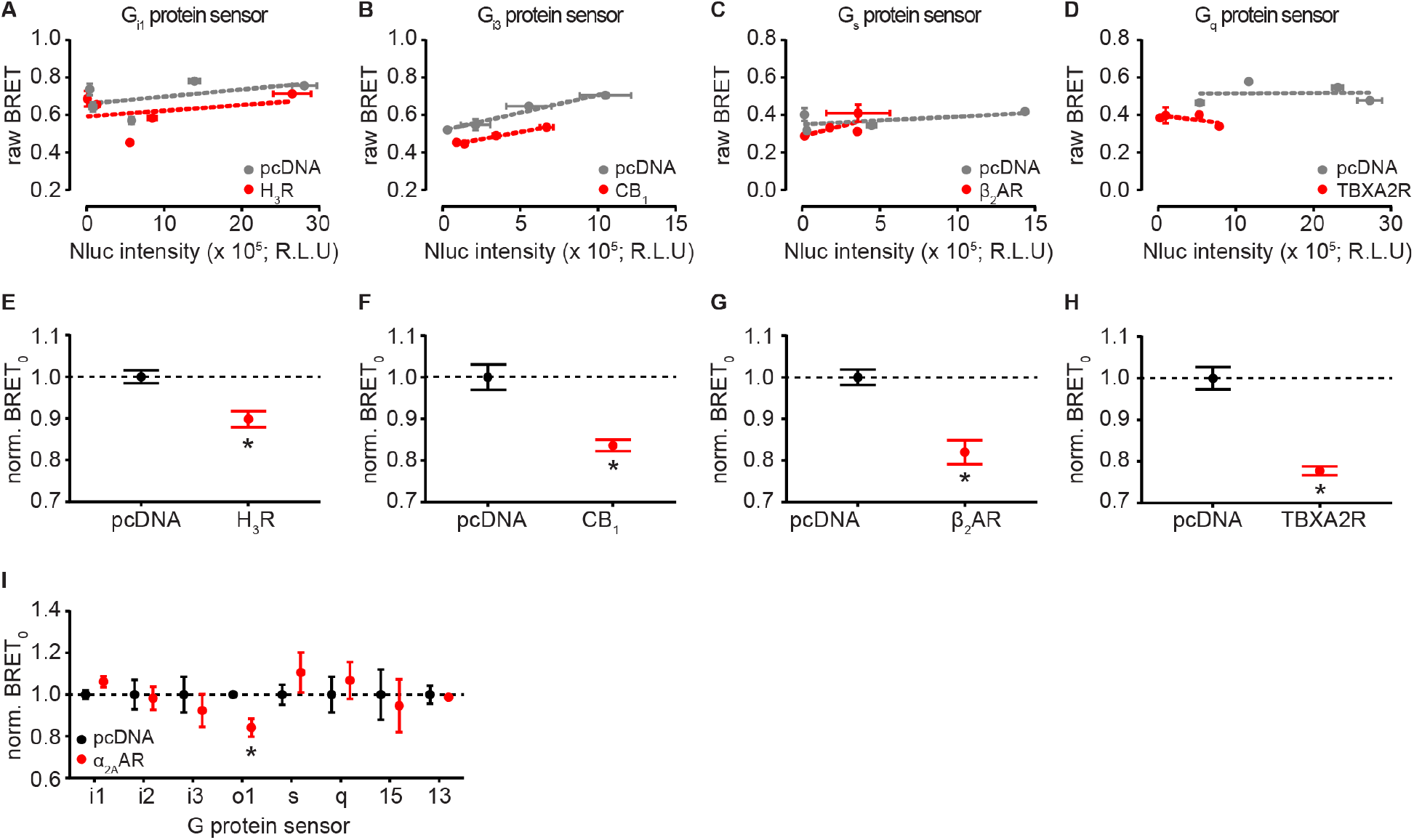
Assessment of constitutive GPCR activity using G protein BRET sensors. **A-D**) BRET over Nluc luminescence plots of HEK293A cells co-transfected with a tricistronic G protein sensor along with pcDNA or a constitutively active GPCR. Data show mean ± s.d. of individual experiments fitted to a linear correlation. **E-H**) Normalized BRET_0_ values ± s.e.m. resulting from the linear fits in (A-D). **I)** Normalized BRET_0_ values of cells co-transfected with the indicated G protein BRET sensor along with either pcDNA or the the α_2A_-adrenergic receptor (α_2A_AR). Statistical difference to pcDNA was tested using Student’s unpaired t-test (p < 0.05). Data show mean ± s.d. (A-D) or s.e.m (E-I) of three to four independent experiments conducted in transiently transfected HEK293A cells.

Next, to employ and validate our approach with another set of constitutively active receptors, we investigated the G protein activity profile of a cluster of orphan class A GPCRs: GPR3, GPR6 and GPR12 (*65*). Expression of these receptors elevates the basal levels of cAMP (*20, 66, 67*), arguing for constitutive signaling through G_s_ and adenylyl cyclases (ACs). In addition, GPR6 and GPR12 have indirectly (i.e. through pharmacological G_i/o_ inhibition) been linked to constitutive activation of G_i/o_ family proteins (*68*). However, evidence for direct activation of G_s_ and G_i/o_ proteins by GPR3/6/12, as well as a comprehensive evaluation of (constitutive) coupling of these orphan GPCRs to other G protein families is currently lacking. By determining the luminescence/BRET correlation of each control and GPR3/6 or 12-transfected sample (**Fig. S6**), we assessed constitutive coupling of these orphan GPCRs to eight distinct G proteins (**Fig. 7A-H**). These data confirmed the constitutive activation of G_s_ by all three orphan GPCRs (**Fig. 7e**), leading to the previously reported induction of the AC/cAMP pathway. Furthermore, we found constitutive activation of G_o1_ by GPR12 (**Fig. 7D**) but no basal signaling activity in any other GPR/G protein pair. In contrast to these results, comparison of raw BRET values without correction for distinct luminescence intensities failed to reveal the constitutive activity of G_s_ and revealed several putative false-positive GPR/G protein pairs (**Fig. S7**).

**Fig. 7:**
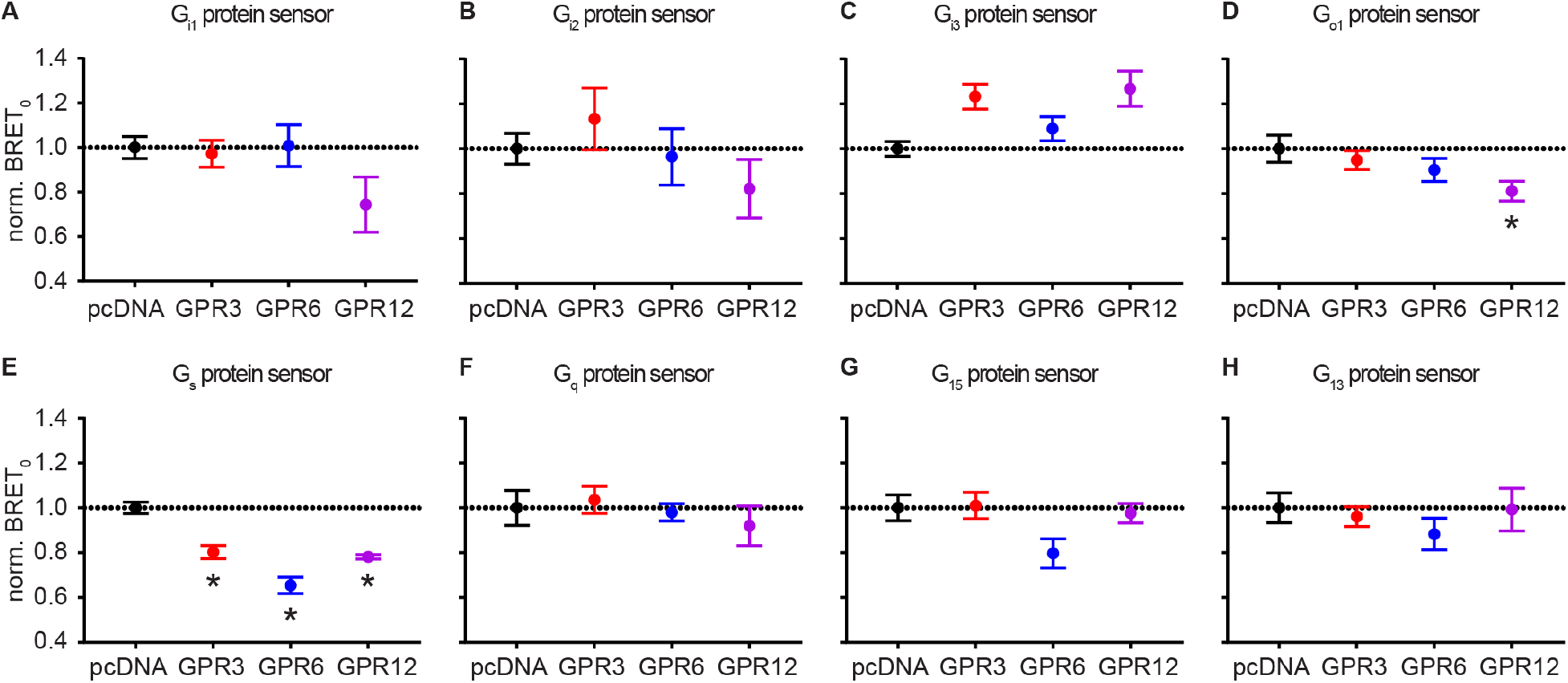
Constitutive G protein activity profiles of the orphan class A GPCRs GPR3, GPR6 and GPR12. Normalized BRET_0_ values ± s.e.m. resulting from the linear fits of BRET over Nluc luminescence plots in Fig. S5 (five to six independent experiments). Statistical difference to pcDNA was tested using One-way ANOVA followed by Dunnett’s multiple comparison (p < 0.05). All experiments were conducted in transiently transfected HEK293A cells.

After confirming that our approach reveals constitutive activity of wildtype GPCRs, we ultimately aimed to screen the activity profile of disease-related receptor mutants. As a model, we chose the class B parathyroid hormone receptor 1 (PTHR1) since this receptor activates both G_s_ and G_q/11_ family members upon binding of its endogenous agonist parathyroid hormone (PTH) (*42*). Furthermore, biochemical studies in the late 90s linked three distinct Jansen-type metaphyseal chondrodysplasia-related PTHR1 point mutants (H^2.50^R, T^6.42^P and I^7.56^R) to enhanced basal levels of cAMP but not of IP_1/2/3_ (*69–72*). We introduced these point mutations into N-terminally HA-tagged PTHR1 and confirmed the functional expression of these receptors at the cell surface (**Fig. S8**). Surprisingly, the G protein BRET experiments with wildtype and mutated PTHR1 indicated a rather promiscuous constitutive signaling profile of the three receptor point mutants (**Fig. 8**, **Fig. S9**). Besides constitutive activation of G_s_ by H^2.50^R and I^7.56^R (but not T^6.42^P), all three mutants also exhibited ligand-independent signaling through the other three major G protein families.

**Fig. 8:**
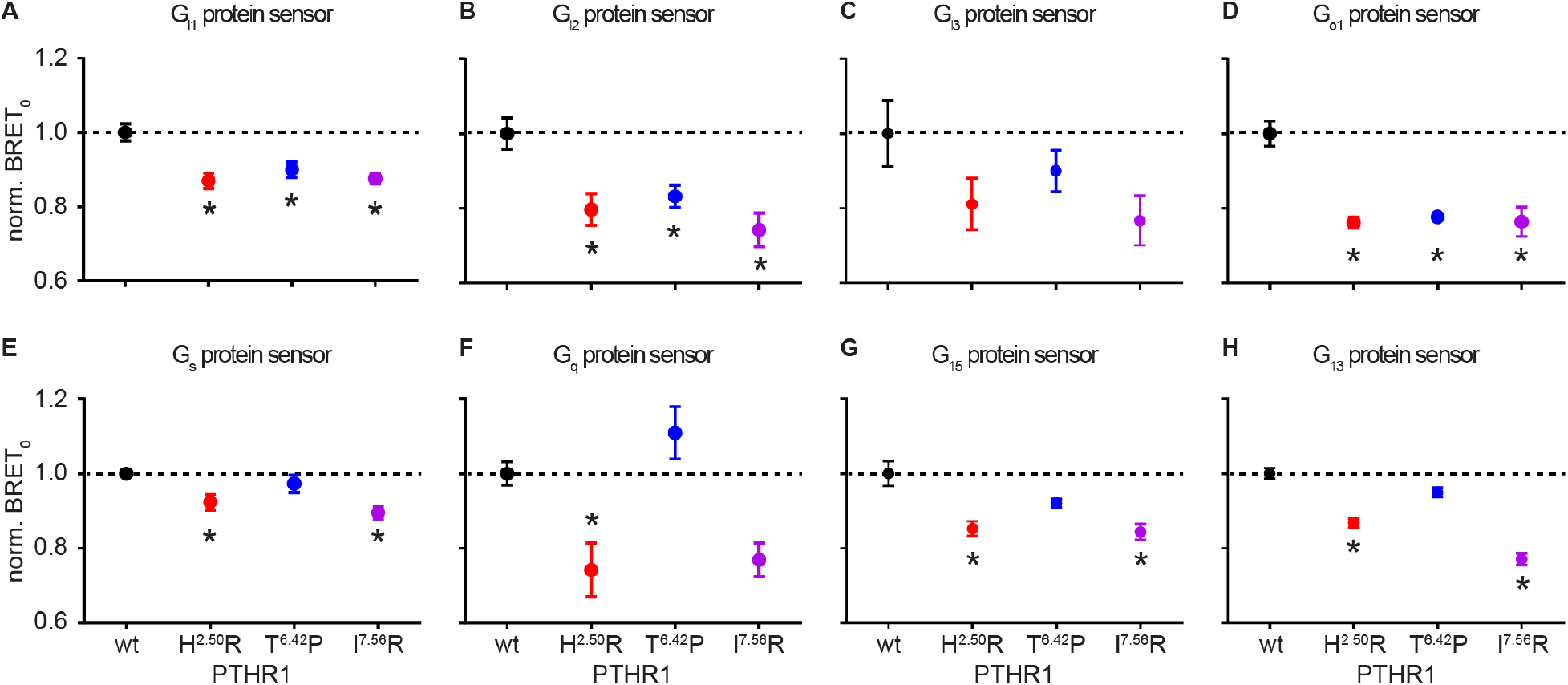
Constitutive G protein activity profiles of Jansen’s disease-related PTHR1 mutants. Normalized BRET_0_ values ± s.e.m. resulting from the linear fits of BRET over Nluc luminescence plots in Fig. S8 (three independent experiments). Statistical difference to wildtype PTHR1 was tested using one-way ANOVA followed by Dunnett’s multiple comparison (p < 0.05). All experiments were conducted in transiently transfected HEK293A cells.

## Discussion

GPCR signaling through heterotrimeric G proteins constitutes the central event in a plethora of physiological and pathological processes. A number of native and disease-related mutant GPCRs exert tissue- and expression level-dependent constitutive activity resulting in ligand-independent activation of G protein-mediated signaling cascades. Due to a lack of sensitive, robust and generalizable techniques to assess GPCR/G protein activity at a proximal level, most studies on constitutive GPCR activity rely on the quantification of second messengers which, themselves, are involved in various (including GPCR- and G protein-independent) cellular pathways, hampering their direct correlation to basal receptor activity.

Here, we addressed this limitation by validating a new cell-based approach relying on optical G protein biosensors that show higher degrees of activity upon co-expression of constitutively active GPCRs. These sensors were obtained by fine-tuning the sensitivity and robustness of previously described FRET- and BRET-based G protein biosensors and by encoding all three sensor subunits on a single tricistronic plasmid. This design resulted in a substantially improved assay sensitivity, which allowed us to establish a new experimental approach to investigate constitutive activity of GPCRs. Furthermore, this plasmid design simplifies the generation of stably expressing cell lines for the purpose of compound screening and hit validation.

The fidelity of our approach to assess constitutive GPCR/G protein activity relies on the correction for varying sensor emission intensities (which artificially affect the BRET ratio, **Fig. 5**) by comparing BRET values at maximum receptor excess (‘BRET_0_’), derived from linear regression of total luminescence and corresponding BRET ratios (**Fig. 6**). This normalization procedure can also be used to correct BRET or FRET signals obtained with other signaling readouts. These include, for instance, the direct assessment of GPCR/G_α_ coupling (*73*) or GPCR/β-arrestin interaction and signaling (*74, 75*), the relative quantification of intracellular second messenger levels, or the measurement of pathway activity in GPCR-dependent bystander BRET assays (*76*). However, the approach is limited to compare only data collected with the very same reading parameters since technical settings (e.g. emission channel widths, luminescence integration time, pre-set gain multiplication factors or the color of the microtiter plates) affect the total number of detected photons and their conversion into electrical currents, thereby shaping the best-fit parameters of linear luminescence/BRET correlations.

On the other hand, the need to overexpress engineered G proteins along with the receptor of interest limits the reliability of our approach as physiologically irrelevant receptor/effector expression ratios can be obtained. Stable expression of these G protein biosensors at physiological levels would benefit the fidelity of this method and labelling of endogenous G protein subunits by gene-editing technologies could enable translation of these findings to in-vivo model systems. Furthermore, it is important to note that these biosensors report on the spatial separation of G_α_ and G_βγ_ subunits but do not directly reveal GDP/GTP exchange in G_α_ subunits as obtained with G_α_-GTP-targeted BRET reporters (*77*). Likewise, unproductive coupling of receptors to heterotrimeric G proteins is not detected by these optical tools as these pairings, by definition, do not result in G protein activation (*78*).

With the considerations discussed above, our sensor toolbox combined with the new analysis scheme provides an advanced biophysical platform to screen for constitutively active GPCRs in living cells with subtype-specific resolution. In contrast to previously employed techniques used to investigate basal GPCR/G protein activity, no application of radiochemical tracers (e.g. [^35^S]GTPγS) or cell lysis is required, and the readout is less amplified or prone to distortion by signaling crosstalk as compared to second messenger-based technologies (e.g. cAMP and IP_1/2/3_).

From a biological perspective, our data obtained with the orphan class A GPCRs and disease-related PTHR1 mutants might be of high relevance for future (orphan) GPCR-directed screening campaigns and for the development of PTHR1-targeting, personalized medication. While our results resembled previous reports on constitutive signaling of GPR3, GPR6 and GPR12 through G_s_ and G_o1_, some of our PTHR1 results contrast the outcome of earlier biochemical investigations. For instance, PTHR1 T^6.42^P did not mediate constitutive G_s_ activity in our setup (in HEK293 cells) despite (partially) elevating cAMP in COS-7 cells (*69, 72*). Furthermore, the significant BRET_0_ signals of PTHR1 H^2.50^R/G_q_, PTHR1 H^2.50^R/G_15_ and PTHR1 I^7.56^R/G_15_ are not consistent with the lack of IP_1/2/3_ elevation (*70–72*). These discrepancies can be due to diminished signal amplification in our G protein-based assay (compared to second-messenger quantification assays) or reflect the distinct G protein and downstream effector (such as adenylyl cyclase II activated by G_βγ_ complexes) expression profiles in HEK293 vs. COS-7 cells.

In summary, the application of this technology and its further adjustment for use in physiologically more relevant samples will help elucidating the basal signaling profiles of GPCRs and aid in a deeper understanding of GPCR/G protein function in health and disease.

## Materials and Methods

### Plasmids and molecular cloning

Plasmids encoding N-terminally FLAG-tagged β_2_AR and HA-tagged PTHR1 were described previously (*39*) and used for the generation of HA-tagged PTHR1 point mutants H^2.50^R, T^6.42^P and I^7.56^R using the Geneart Site-directed Mutagenesis Kit (Thermo Fisher Scientific). Plasmid encoding wild-type TBXA2R was kindly provided by A. Inoue (Tohoku University, Sendai, Japan). Wildtype H_3_R, GPR3, GPR6, GPR12 and G protein subunits were either purchased from cDNA.org in pcDNA vectors or amplified from previously described G protein sensors (*34*). PcDNA-G_αo1_-Nluc, pcDNA-G_α13_-Nluc and pcDNA-G_α15_-Nluc were synthesized by GenScript. The subunits were then fused to cpVenus^173^ or Nluc at the indicated positions (**Fig. 1c**) and encoded on a tricistronic T2A-IRES vector as described previously (*32,34*) using established PCR techniques and restriction enzymes. All constructs were verified by sequencing (Eurofins genomics).

### Reagents

Histamine dihydrochloride was purchased from Tocris Bioscience (Wiesbaden-Nordenstadt, Germany). Poly-D-lysine (PDL), isoprenaline, ICI118.551 and U-46619 were obtained from Sigma Aldrich (Merck KGaA, Darmstadt, Germany). The Nluc substrate furimazine was from Promega (Madison, WI, USA) and white-wall, white-bottomed 96-well, as well as flat-bottomed transparent microtiter plates were from Gibco (Waltham, MA, USA).

### Cell culture

HEK293A cells were used for transient expression of GPCRs and G protein biosensors and grown in Dulbecco’s Modified Eagle’s Medium (DMEM) supplemented with 2 mM glutamine, 10% fetal calf serum, 0.1 mg/mL streptomycin, and 100 units/mL penicillin at 37 °C with 5% CO_2_.

### Transient transfection and plating

Resuspended cells (300,000 cells / mL) were transfected in suspension with a total of 1 μg DNA / mL suspension using Lipofectamine 2000 (Thermo Fisher Scientific, Waltham, MA, USA). For transfection of PTHR1 constructs only, 1 μg of the plasmids were transfected per mL cell suspension. Titration experiments shown in **Fig. 5** were obtained by transfecting 1-100% sensor DNA in a total of 1 μg DNA per mL cell suspension (added up with pcDNA). For co-transfection of GPCRs along with tricistronic G protein sensors, 500 ng GPCR and 500 ng G protein sensor were combined. To transfect TBXA2R along with plasmids separately encoding the G_13_ sensor subunits, 500 ng GPCR plasmid were combined with 167 ng pcDNA-G_α13_-Nluc, 167 ng pcDNA-G_β3_ and 167 ng pcDNA-cpVenus^173^-G_γ9_. Cells mixed with the transfection reagents were seeded onto PDL-pre-coated 96-well plates and grown for 48 hours at 37 °C with 5% CO_2_. White plates were used for the recording of luminescence spectra and BRET experiments, transparent, flat bottom 96-well plates were used for the assessment of PTHR1 surface levels.

### Assessment of PTHR1 surface expression through live-cell ELISA

For quantification of cell surface receptor expression, HEK293A cells transfected with pcDNA or N-terminally HA-tagged PTHR1 constructs were grown for 48 hours in transparent 96-well plates and washed once with 0.5% BSA in PBS. Next, cells were incubated with a rabbit anti-HA-tag antibody (1 μg/ml, cat# ab9110; Abcam) in 1% BSA–PBS for 1 h at 4 °C. Following incubation, the cells were washed three times with 0.5% BSA–PBS and incubated with a horseradish peroxidase-conjugated goat anti-rabbit antibody (0.3 μg/ml, cat# 31460; Thermo Fisher Scientific) in 1% BSA–PBS for 1 h at 4 °C. The cells were washed three times with 0.5% BSA/PBS, and 50 μl of the peroxidase substrate 3,3′,5,5′-tetramethylbenzidine (T8665; Sigma-Aldrich) was added. Subsequently, the cells were incubated for 20 min and 50 μl of 2 M HCl was added. The absorbance was read at 450 nm using a BMG Ω POLARstar plate reader.

### Recording of G protein luminescence spectra

HEK293A cells were transfected and seeded as described above. Luminescence emission of the G protein sensors was recorded between 400 nm and 600 nm with 2 nm resolution in HBSS upon addition of 1:1,000 furimazine dilution. All experiments were conducted using a CLARIOstar plate reader (BMG, Ortenberg, Germany) and spectra were normalized to the donor emission peak.

### BRET measurements

Transfected cells grown for 48 hours in 96-well plates were washed with HBSS and incubated with 1/1,000 dilution of furimazine stock solution. After incubation for 3 minutes at 37°C, the BRET ratio was measured in two consecutive reads for the assessment of constitutive receptor activity. To assess GPCR ligand-induced changes in G protein BRET, three basal reads were followed by addition of ligand solutions or vehicle control and subsequent BRET reads for to detect ligand-induced changes in BRET. All experiments were conducted at 37 °C with a CLARIOstar plate reader. Nluc emission intensity was selected using a 450/40 nm monochromator (Gain: 3600) and cpVenus^173^ emission using a 535/30 nm monochromator (Gain: 4000) with an integration time of 0.3 seconds in both channels.

### Data analysis

BRET ratios were defined as acceptor emission/donor emission. The basal BRET ratio before ligand stimulation (Ratio_basal_) was defined as the average of three consecutive BRET values (respectively nine consecutive basal reads for the data presented in Fig. S5). To quantify ligand-induced changes, ΔBRET was calculated for each well as a percent over basal [(Ratio_stim_− Ratio_basal_)/ Ratio_basal_] × 100). Subsequently, the average ΔBRET of vehicle control was subtracted. Three consecutive BRET ratios (Ratio_stim_) were averaged for the generation of ΔBRET ligand concentration-response curves and Z-factor analyses. Z-factors were calculated according to Zhang et al. (*44*) based on the following equation:

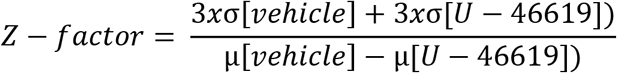

where σ and μ are the standard deviations and average ΔBRET values of 10 μM U-46619 and vehicle control, respectively.

Data were analysed using Prism 5.0 software (GraphPad, San Diego, CA, USA). Data from BRET and luminescence concentration–response experiments were fitted using a four-parameter fit. Nluc/BRET plots were fitted using a linear fit and tested for deviation from linear correlation applying runs test (p < 0.05). BRET_0_ was defined as the Y-intercept with its computed standard error resulting from the linear fit of BRET values over increasing Nluc intensities. Differences between BRET_0_ values were tested for significance using either Student’s t-test (**Fig. 6**) or one-way ANOVA followed by Dunnett’s multiple comparison (**Fig. 7, 8**) as indicated in the figure legends. P values < 0.05 were considered significant.

## Supplementary Materials

Fig. S1: Bioluminescence spectra of tricistronic G protein biosensors.

Fig. S2: Z-factor data used to assess assay improvement with the tricistronic sensor design.

Fig. S3: Assessment of constitutive GPCR activity without correction for distinct G protein sensor emission intensities.

Fig. S4: BRET over Nluc plots for the assessment of constitutive G protein activity profiles in the orphan class A GPCR cluster GPR3, GPR6 and GPR12.

Fig. S5: Effect of the inverse agonist rimonabant on CB_1_ receptor-mediated G_i3_ sensor activation.

Fig. S6: BRET over Nluc plots for the assessment of constitutive G protein activity profiles in the orphan class A GPCR cluster GPR3, GPR6 and GPR12.

Fig. S7: Assessment of constitutive orphan GPCR activity without correction for distinct G protein sensor emission intensities.

Fig. S8: Surface expression levels of PTHR1 point mutants.

Fig. S9: BRET over Nluc plots for the assessment of constitutive G protein activity profiles orphan Jansen’s disease-related PTHR1 mutants.

Table S1. P-values of runs test for deviation from linear correlation in Figure 5.

## Acknowledgments

We thank Anna Krook at the Dept. Physiology & Pharmacology, Karolinska Institutet for providing access to the CLARIOstar plate reader; Asuka Inoue at the Tohoku University, Sendai for the plasmid encoding TBXA2R; Ulrike Zabel at the Institute of Pharmacology and Toxicology, University of Wuerzburg for cloning support and Nevin Lambert at Augusta University, Georgia for helpful discussions.

## Author contributions

H.S. and G.S. conceived, designed and coordinated the study. H.S. and R.S. performed experiments. H.S. prepared the figures with input from all authors. H.S. wrote the manuscript and all authors revised the manuscript. G.S. supervised and financed the project.

## Data and materials availability

The raw data are available upon request to the corresponding authors. All eight tricistronic G protein sensors are available via the plasmid repository www.addgene.org.

## References and Notes

1. K. Sriram, P. A. Insel, G Protein-Coupled Receptors as Targets for Approved Drugs: How Many Targets and How Many Drugs? Mol Pharmacol 93, 251–258 (2018).

2. J. R. Hepler, A. G. Gilman, G proteins. Trends Biochem Sci 17, 383–387 (1992).

3. K. Y. Chung et al., Conformational changes in the G protein Gs induced by the beta2 adrenergic receptor. Nature 477, 611–615 (2011).

4. R. O. Dror et al., SIGNAL TRANSDUCTION. Structural basis for nucleotide exchange in heterotrimeric G proteins. Science 348, 1361–1365 (2015).

5. M. Bunemann, M. Frank, M. J. Lohse, Gi protein activation in intact cells involves subunit rearrangement rather than dissociation. Proc Natl Acad Sci U S A 100, 16077–16082 (2003).

6. M. Frank, L. Thumer, M. J. Lohse, M. Bunemann, G Protein activation without subunit dissociation depends on a G{alpha}(i)-specific region. J Biol Chem 280, 24584–24590 (2005).

7. G. J. Digby, R. M. Lober, P. R. Sethi, N. A. Lambert, Some G protein heterotrimers physically dissociate in living cells. Proc Natl Acad Sci U S A 103, 17789–17794 (2006).

8. C. Gales et al., Probing the activation-promoted structural rearrangements in preassembled receptor-G protein complexes. Nat Struct Mol Biol 13, 778–786 (2006).

9. J. R. Hepler, G protein coupled receptor signaling complexes in live cells. Cell Logist 4, e29392 (2014).

10. N. A. Lambert, Dissociation of heterotrimeric g proteins in cells. Sci Signal 1, re5 (2008).

11. Y. K. Chung, Y. H. Wong, Re-examining the ‘Dissociation Model’ of G protein activation from the perspective of Gbetagamma signaling. FEBS J, (2020).

12. M. Fukami, E. Suzuki, M. Igarashi, M. Miyado, T. Ogata, Gain-of-function mutations in G-protein-coupled receptor genes associated with human endocrine disorders. Clin Endocrinol (Oxf) 88, 351–359 (2018).

13. R. Seifert, K. Wenzel-Seifert, Constitutive activity of G-protein-coupled receptors: cause of disease and common property of wild-type receptors. Naunyn Schmiedebergs Arch Pharmacol 366, 381–416 (2002).

14. X. Han, Constitutively active chemokine CXC receptors. Adv Pharmacol 70, 265–301 (2014).

15. F. J. Meye, G. M. Ramakers, R. A. Adan, The vital role of constitutive GPCR activity in the mesolimbic dopamine system. Transl Psychiatry 4, e361 (2014).

16. M. J. Smit et al., Pharmacogenomic and structural analysis of constitutive g protein-coupled receptor activity. Annu Rev Pharmacol Toxicol 47, 53–87 (2007).

17. T. Costa, A. Herz, Antagonists with negative intrinsic activity at delta opioid receptors coupled to GTP-binding proteins. Proc Natl Acad Sci U S A 86, 7321–7325 (1989).

18. M. A. Kjelsberg, S. Cotecchia, J. Ostrowski, M. G. Caron, R. J. Lefkowitz, Constitutive activation of the alpha 1B-adrenergic receptor by all amino acid substitutions at a single site. Evidence for a region which constrains receptor activation. J Biol Chem 267, 1430–1433 (1992).

19. P. Chidiac, T. E. Hebert, M. Valiquette, M. Dennis, M. Bouvier, Inverse agonist activity of beta-adrenergic antagonists. Mol Pharmacol 45, 490–499 (1994).

20. D. Eggerickx et al., Molecular cloning of an orphan G-protein-coupled receptor that constitutively activates adenylate cyclase. Biochem J 309 (Pt 3), 837–843 (1995).

21. G. Barreda-Gomez, M. Teresa Giralt, R. Rodriguez-Puertas, Methods to measure g-protein-coupled receptor activity for the identification of inverse agonists. Methods Enzymol 485, 261–273 (2010).

22. S. E. Wu, W. E. Miller, The HCMV US28 vGPCR induces potent Galphaq/PLC-beta signaling in monocytes leading to increased adhesion to endothelial cells. Virology 497, 233–243 (2016).

23. T. Muroi et al., GPR62 constitutively activates cAMP signaling but is dispensable for male fertility in mice. Reproduction 154, 755–764 (2017).

24. M. L. J. Doornbos et al., Constitutive activity of the metabotropic glutamate receptor 2 explored with a whole-cell label-free biosensor. Biochem Pharmacol 152, 201–210 (2018).

25. Q. Wang, X. Dong, J. Lu, T. Hu, G. Pei, Constitutive activity of a G protein-coupled receptor, DRD1, contributes to human cerebral organoid formation. Stem Cells 38, 653–665 (2020).

26. R. Link et al., The constitutive activity of melanocortin-4 receptors in cAMP pathway is allosterically modulated by zinc and copper ions. J Neurochem 153, 346–361 (2020).

27. R. L. Beckner, L. Zoubak, K. G. Hines, K. Gawrisch, A. A. Yeliseev, Probing thermostability of detergent-solubilized CB2 receptor by parallel G protein-activation and ligand-binding assays. J Biol Chem 295, 181–190 (2020).

28. R. H. J. Olsen et al., TRUPATH, an open-source biosensor platform for interrogating the GPCR transducerome. Nat Chem Biol 16, 841–849 (2020).

29. M. J. Adjobo-Hermans et al., Real-time visualization of heterotrimeric G protein G_q_ activation in living cells. BMC Biol 9, 32 (2011).

30. M. Busnelli et al., Functional selective oxytocin-derived agonists discriminate between individual G protein family subtypes. J Biol Chem 287, 3617–3629 (2012).

31. C. Janetopoulos, T. Jin, P. Devreotes, Receptor-mediated activation of heterotrimeric G-proteins in living cells. Science 291, 2408–2411 (2001).

32. M. Mastop et al., A FRET-based biosensor for measuring Galpha13 activation in single cells. PLoS One 13, e0193705 (2018).

33. A. Sauliere et al., Deciphering biased-agonism complexity reveals a new active AT1 receptor entity. Nat Chem Biol 8, 622–630 (2012).

34. J. van Unen et al., A New Generation of FRET Sensors for Robust Measurement of Galphai1, Galphai2 and Galphai3 Activation Kinetics in Single Cells. PLoS One 11, e0146789 (2016).

35. H. Yano et al., Development of novel biosensors to study receptor-mediated activation of the G-protein alpha subunits G_s_ and Golf. J Biol Chem 292, 19989–19998 (2017).

36. C. Gales et al., Real-time monitoring of receptor and G-protein interactions in living cells. Nat Methods 2, 177–184 (2005).

37. J. Z. Yu, M. M. Rasenick, Real-time visualization of a fluorescent G(alpha)(s): dissociation of the activated G protein from plasma membrane. Mol Pharmacol 61, 352–359 (2002).

38. M. P. Hallet al., Engineered luciferase reporter from a deep sea shrimp utilizing a novel imidazopyrazinone substrate. ACS Chem Biol 7, 1848–1857 (2012).

39. H. Schihada et al., A universal bioluminescence resonance energy transfer sensor design enables high-sensitivity screening of GPCR activation dynamics. Commun Biol 1, 105 (2018).

40. J. Klarenbeek, J. Goedhart, A. van Batenburg, D. Groenewald, K. Jalink, Fourth-generation epac-based FRET sensors for cAMP feature exceptional brightness, photostability and dynamic range: characterization of dedicated sensors for FLIM, for ratiometry and with high affinity. PLoS One 10, e0122513 (2015).

41. A. L. Szymczak, D. A. Vignali, Development of 2A peptide-based strategies in the design of multicistronic vectors. Expert Opin Biol Ther 5, 627–638 (2005).

42. A. Inoue et al., Illuminating G-Protein-Coupling Selectivity of GPCRs. Cell 177, 1933–1947 e1925 (2019).

43. H. Schihada et al., Development of a conformational histamine H3 receptor biosensor for the synchronous screening of agonists and inverse agonists. ACS Sens, (2020).

44. J. H. Zhang, T. D. Chung, K. R. Oldenburg, A Simple Statistical Parameter for Use in Evaluation and Validation of High Throughput Screening Assays. J Biomol Screen 4, 67–73 (1999).

45. Y. Wang, P. D. Townsend, Common mistakes in luminescence analysis. J Phys Conf Ser 398, (2012).

46. M. Sato, J. B. Blumer, V. Simon, S. M. Lanier, Accessory proteins for G proteins: partners in signaling. Annu Rev Pharmacol Toxicol 46, 151–187 (2006).

47. E. M. Ross, T. M. Wilkie, GTPase-activating proteins for heterotrimeric G proteins: regulators of G protein signaling (RGS) and RGS-like proteins. Annu Rev Biochem 69, 795–827 (2000).

48. K. Wieland et al., Constitutive activity of histamine h(3) receptors stably expressed in SK-N-MC cells: display of agonism and inverse agonism by H(3) antagonists. J Pharmacol Exp Ther 299, 908–914 (2001).

49. J. M. Arrang, S. Morisset, F. Gbahou, Constitutive activity of the histamine H3 receptor. Trends Pharmacol Sci 28, 350–357 (2007).

50. A. Rouleau et al., Histamine H3-receptor-mediated [35S]GTP gamma[S] binding: evidence for constitutive activity of the recombinant and native rat and human H3 receptors. Br J Pharmacol 135, 383–392 (2002).

51. D. Moreno-Delgado et al., Constitutive activity of H3 autoreceptors modulates histamine synthesis in rat brain through the cAMP/PKA pathway. Neuropharmacology 51, 517–523 (2006).

52. S. Morisset et al., High constitutive activity of native H3 receptors regulates histamine neurons in brain. Nature 408, 860–864 (2000).

53. C. C. Felder et al., Comparison of the pharmacology and signal transduction of the human cannabinoid CB1 and CB2 receptors. Mol Pharmacol 48, 443–450 (1995).

54. M. Canals, G. Milligan, Constitutive activity of the cannabinoid CB1 receptor regulates the function of co-expressed Mu opioid receptors. J Biol Chem 283, 11424–11434 (2008).

55. B. Fioravanti et al., Constitutive activity at the cannabinoid CB1 receptor is required for behavioral response to noxious chemical stimulation of TRPV1: antinociceptive actions of CB1 inverse agonists. J Neurosci 28, 11593–11602 (2008).

56. J. Nie, D. L. Lewis, Structural domains of the CB1 cannabinoid receptor that contribute to constitutive activity and G-protein sequestration. J Neurosci 21, 8758–8764 (2001).

57. A. Lattion, L. Abuin, M. Nenniger-Tosato, S. Cotecchia, Constitutively active mutants of the beta1-adrenergic receptor. FEBS Lett 457, 302–306 (1999).

58. S. Engelhardt, Y. Grimmer, G. H. Fan, M. J. Lohse, Constitutive activity of the human beta(1)-adrenergic receptor in beta(1)-receptor transgenic mice. Mol Pharmacol 60, 712–717 (2001).

59. Y. Y. Zhou et al., Spontaneous activation of beta(2)-but not beta(1)-adrenoceptors expressed in cardiac myocytes from beta(1)beta(2) double knockout mice. Mol Pharmacol 58, 887–894 (2000).

60. H. E. Hopkinson, M. L. Latif, S. J. Hill, Non-competitive antagonism of beta(2)-agonist-mediated cyclic AMP accumulation by ICI 118551 in BC3H1 cells endogenously expressing constitutively active beta(2)-adrenoceptors. Br J Pharmacol 131, 124–130 (2000).

61. A. Chillar, J. Wu, V. Cervantes, K. H. Ruan, Structural and functional analysis of the C-terminus of Galphaq in complex with the human thromboxane A2 receptor provides evidence of constitutive activity. Biochemistry 49, 6365–6374 (2010).

62. R. Chakraborty et al., Site-directed mutations and the polymorphic variant Ala160Thr in the human thromboxane receptor uncover a structural role for transmembrane helix 4. PLoS One 7, e29996 (2012).

63. R. Chakraborty, R. P. Bhullar, S. Dakshinamurti, J. Hwa, P. Chelikani, Inverse agonism of SQ 29,548 and Ramatroban on Thromboxane A2 receptor. PLoS One 9, e85937 (2014).

64. P. J. Pauwels, S. Tardif, T. Wurch, F. C. Colpaert, Facilitation of constitutive alpha(2A)-adrenoceptor activity by both single amino acid mutation (Thr(373)Lys) and G(alpha o) protein coexpression: Evidence for inverse agonism. Journal of Pharmacology and Experimental Therapeutics 292, 654–663 (2000).

65. P. Morales, I. Isawi, P. H. Reggio, Towards a better understanding of the cannabinoid-related orphan receptors GPR3, GPR6, and GPR12. Drug Metab Rev 50, 74–93 (2018).

66. K. Uhlenbrock, H. Gassenhuber, E. Kostenis, Sphingosine 1-phosphate is a ligand of the human gpr3, gpr6 and gpr12 family of constitutively active G protein-coupled receptors. Cell Signal 14, 941–953 (2002).

67. J. N. Bresnick et al., Identification of signal transduction pathways used by orphan g protein-coupled receptors. Assay Drug Dev Technol 1, 239–249 (2003).

68. A. L. Martin, M. A. Steurer, R. S. Aronstam, Constitutive Activity among Orphan Class-A G Protein Coupled Receptors. PLoS One 10, e0138463 (2015).

69. T. J. Gardella et al., Inverse agonism of amino-terminally truncated parathyroid hormone (PTH) and PTH-related peptide (PTHrP) analogs revealed with constitutively active mutant PTH/PTHrP receptors. Endocrinology 137, 3936–3941 (1996).

70. E. Schipani, K. Kruse, H. Juppner, A constitutively active mutant PTH-PTHrP receptor in Jansen-type metaphyseal chondrodysplasia. Science 268, 98–100 (1995).

71. E. Schipani et al., A novel parathyroid hormone (PTH)/PTH-related peptide receptor mutation in Jansen’s metaphyseal chondrodysplasia. J Clin Endocrinol Metab 84, 3052–3057 (1999).

72. E. Schipani et al., Constitutively activated receptors for parathyroid hormone and parathyroid hormone-related peptide in Jansen’s metaphyseal chondrodysplasia. N Engl J Med 335, 708–714 (1996).

73. Q. Wan et al., Mini G protein probes for active G protein-coupled receptors (GPCRs) in live cells. J Biol Chem 293, 7466–7473 (2018).

74. F. F. Hamdan, M. Audet, P. Garneau, J. Pelletier, M. Bouvier, High-throughput screening of G protein-coupled receptor antagonists using a bioluminescence resonance energy transfer 1-based beta-arrestin2 recruitment assay. J Biomol Screen 10, 463–475 (2005).

75. A. Oishi, J. Dam, R. Jockers, beta-Arrestin-2 BRET Biosensors Detect Different beta-Arrestin-2 Conformations in Interaction with GPCRs. ACS Sens 5, 57–64 (2020).

76. Y. Namkung et al., Functional selectivity profiling of the angiotensin II type 1 receptor using pathway-wide BRET signaling sensors. Sci Signal 11, (2018).

77. M. Maziarz et al., Revealing the Activity of Trimeric G-proteins in Live Cells with a Versatile Biosensor Design. Cell 182, 770–785 e716 (2020).

78. N. Okashah et al., Agonist-induced formation of unproductive receptor-G12 complexes. Proc Natl Acad Sci U S A 117, 21723–21730 (2020).

