## Supplemental Material for "Quantitative assessment of constitutive G protein-coupled receptor activity with BRET-based G protein biosensors"

### Table of Content

Fig. S1: Bioluminescence spectra of tricistronic G protein biosensors.

Fig. S2: Z-factor data used to assess assay improvement with the tricistronic sensor design.

Fig. S3: Assessment of constitutive GPCR activity without correction for distinct G protein sensor emission intensities.

Fig. S4: BRET over Nluc plots for the assessment of constitutive G protein activity profiles in the orphan class A GPCR cluster GPR3, GPR6 and GPR12.

Fig. S5: Effect of the inverse agonist rimonabant on $CB_1$ receptor-mediated $G_{i3}$ sensor activation.

Fig. S6: BRET over Nluc plots for the assessment of constitutive G protein activity profiles in the orphan class A GPCR cluster GPR3, GPR6 and GPR12.

Fig. S7: Assessment of constitutive orphan GPCR activity without correction for distinct G protein sensor emission intensities.

Fig. S8: Surface expression levels of PTHR1 point mutants.

Fig. S9: BRET over Nluc plots for the assessment of constitutive G protein activity profiles orphan Jansen's disease-related PTHR1 mutants.

Table S1. P-values of runs test for deviation from linear correlation in Figure 5.

**Supplementary Figure S1**

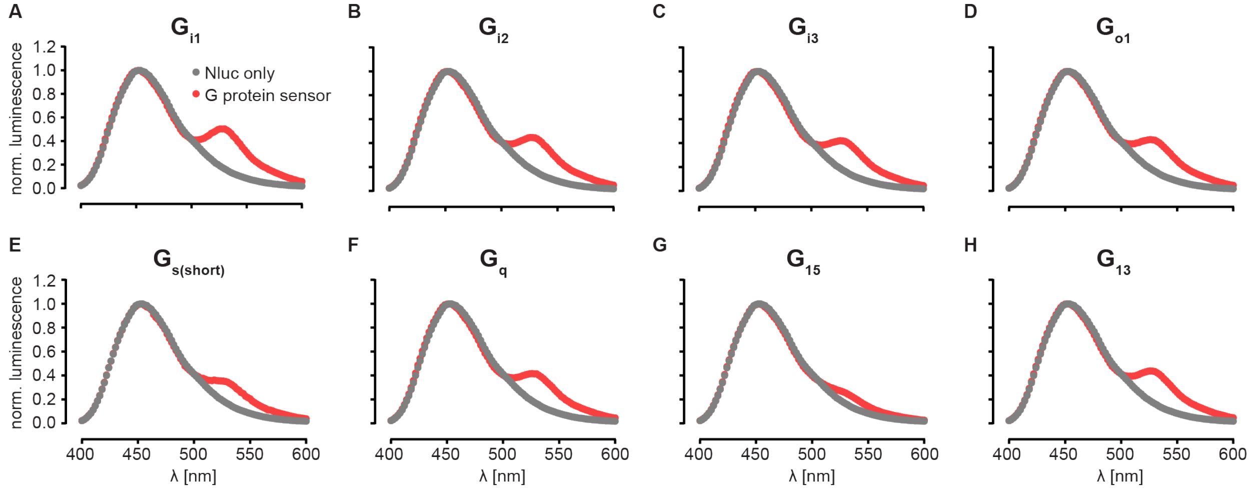


**Fig. S1**: **Bioluminescence spectra of tricistronic G protein biosensors.** The tricistronic plasmids encoding all three G protein subunits were transiently transfected into HEK293A cells. Data show mean ± s.d. of three wells normalized to the donor emission peak.

### Supplementary Figure S2

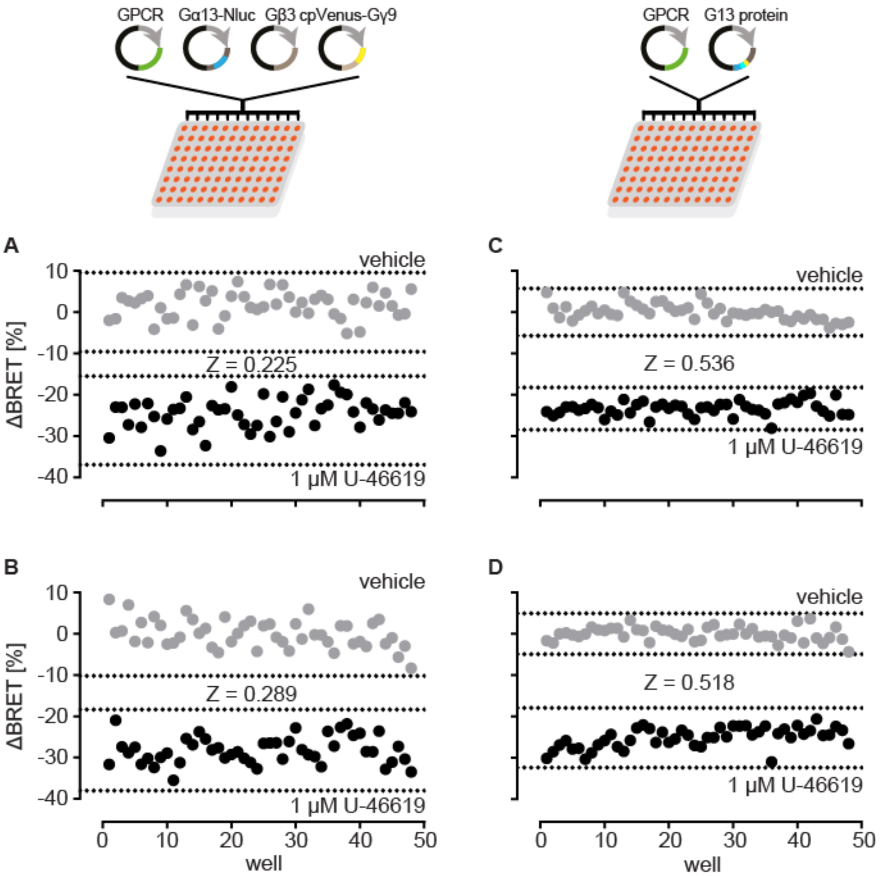


**Fig. S2: Z-factor data used to assess assay improvement with the tricistronic sensor design.** **A, B**) ΔBRET signals of the $G_{13}$ BRET biosensor upon separate co-transfection with the three G protein subunits and TBXA2R. **C, D**) ΔBRET signals of the $G_{13}$ BRET biosensor upon transfection of the tricistronic G protein sensor plasmid along with TBXA2R. The dotted lines represent mean ± three-fold standard deviation of vehicle- and TBXA2R agonist (1 µM U46619)-induced BRET changes. All experiments were conducted in transiently transfected HEK293A cells.

**Supplementary Figure S3**

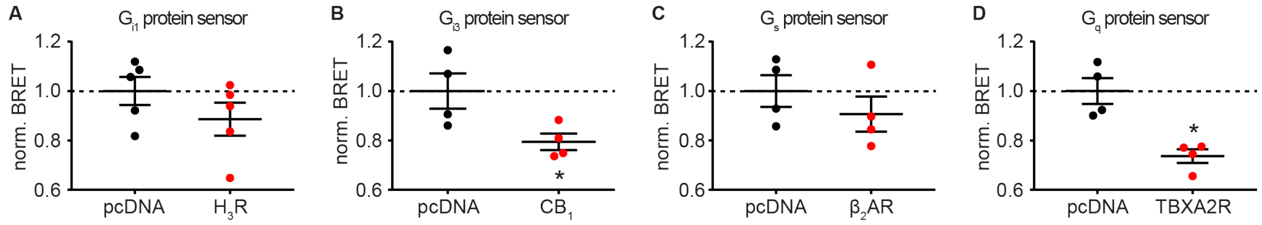


**Fig. S3: Assessment of constitutive GPCR activity without correction for distinct G protein sensor emission intensities.** Normalized BRET values of tricistronic G protein biosensors co-transfected with pcDNA or constitutively active GPCRs into HEK293A cells. The raw BRET values are not corrected for different Nluc emission intensities and normalized to the average BRET value of pcDNA. Data show individual datapoints and mean ± s.e.m. of four to five independent experiments. Statistical difference to pcDNA was assessed using paired Student's t-test ($p < 0.05$).

**Supplementary Figure S4**

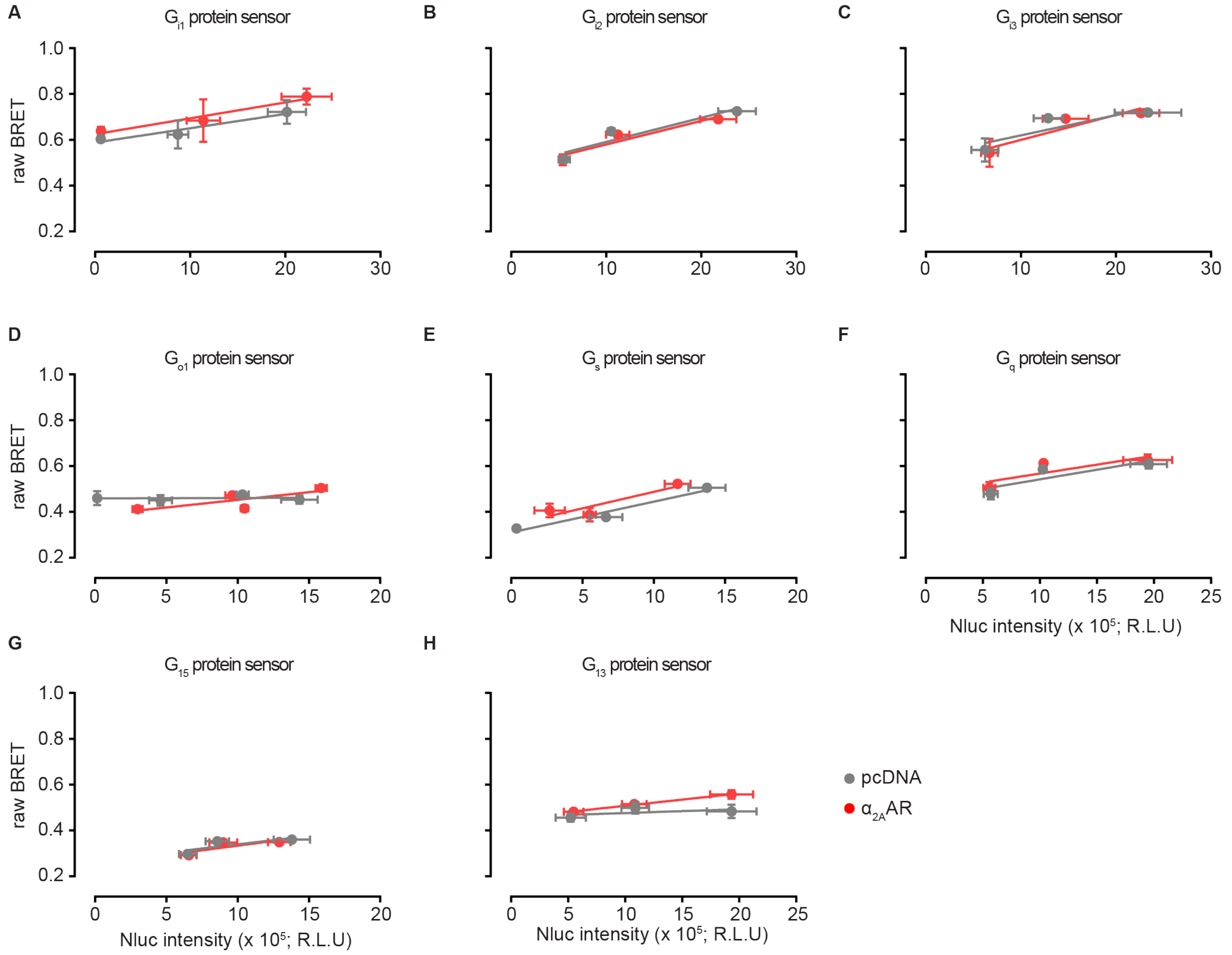


**Fig. S4: BRET over Nluc plots for the assessment of constitutive activity of the $\alpha_{2A}$-adrenergic receptor.** BRET over Nluc luminescence plots of HEK293A cells co-transfected with a tricistronic G protein sensor along with pcDNA or $\alpha_{2A}$AR. Data show mean ± s.d. of three to four individual experiments fitted to a linear correlation.

**Supplementary Figure S5**

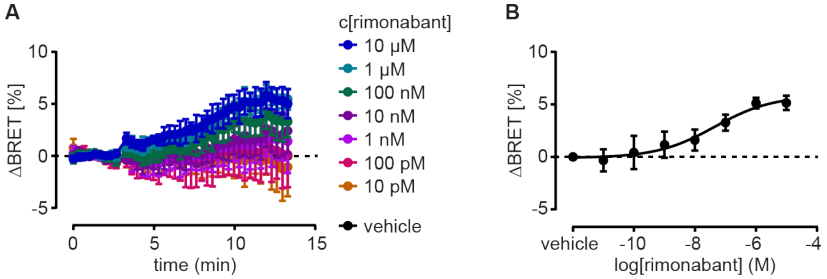


**Fig. S5: Effect of the inverse agonist rimonabant on $CB_1$ receptor-mediated $G_{i3}$ sensor activation. A)** ΔBRET time-course induced by the $CB_1$ receptor inverse agonist rimonabant. **B)** ΔBRET concentration-response curve of rimonabant after 8 minutes incubation. All experiments were conducted in HEK293A cells co-transfected with $CB_1$ and the tricistronic $G_{i3}$ protein sensor. Data show mean ± s.e.m. of three individual experiments.

**Supplementary Figure S6**

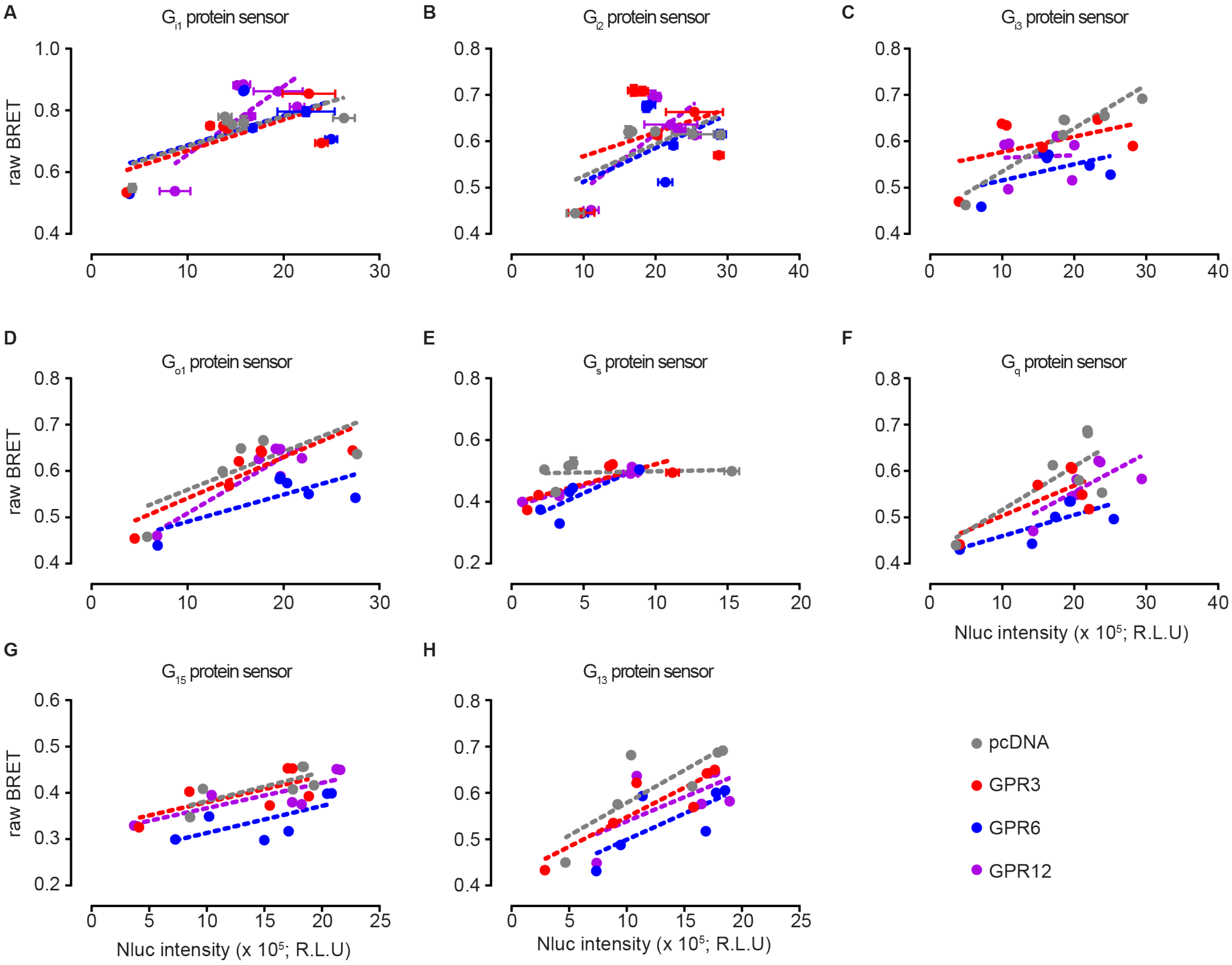


**Fig. S6: BRET over Nluc plots for the assessment of constitutive G protein activity profiles in the orphan class A GPCR cluster GPR3, GPR6 and GPR12.** BRET over Nluc luminescence plots of HEK293A cells co-transfected with a tricistronic G protein sensor along with pcDNA or the indicated orphan GPCR. Data show mean ± s.d. of five to six individual experiments fitted to a linear correlation.

**Supplementary Figure S7**

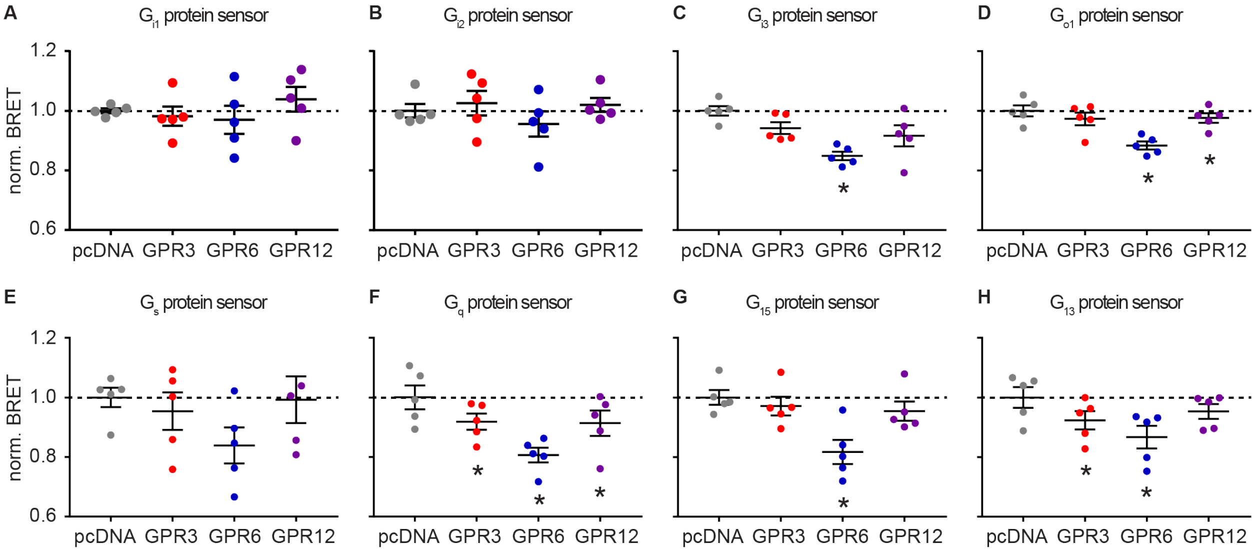


**Fig. S7: Assessment of constitutive orphan GPCR activity without correction for distinct G protein sensor emission intensities.** Normalized BRET values of tricistronic G protein biosensors co-transfected with pcDNA or one orphan class A GPCR into HEK293A cells. The raw BRET values were not corrected for different Nluc emission intensities and normalized to the average BRET value of pcDNA. Data show individual datapoints and mean ± s.e.m. of five to six independent experiments. Statistical difference to pcDNA was assessed using paired one-way ANOVA followed by Dunnett's multiple comparison ($p < 0.05$).

**Supplementary Figure S8**

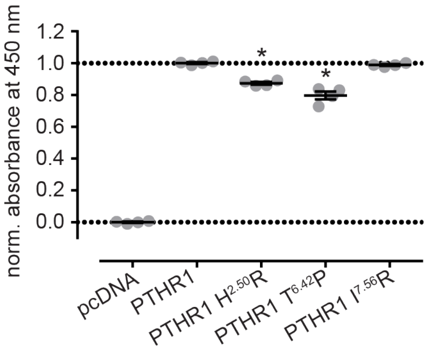


**Fig. S8: Surface expression levels of PTHR1 point mutants.** Cell surface ELISA experiments were conducted in transiently transfected HEK293A cells. Empty vector (pcDNA)-transfected wells were used for background subtraction and all values were normalized to the average absorbance of PTHR1 wildtype. Data show mean ± s.e.m. of four independent experiments and statistical difference to wildtype PTHR1 was assessed through one-way ANOVA followed by Dunnett's multiple comparison ($p < 0.05$).

**Supplementary Figure S9**

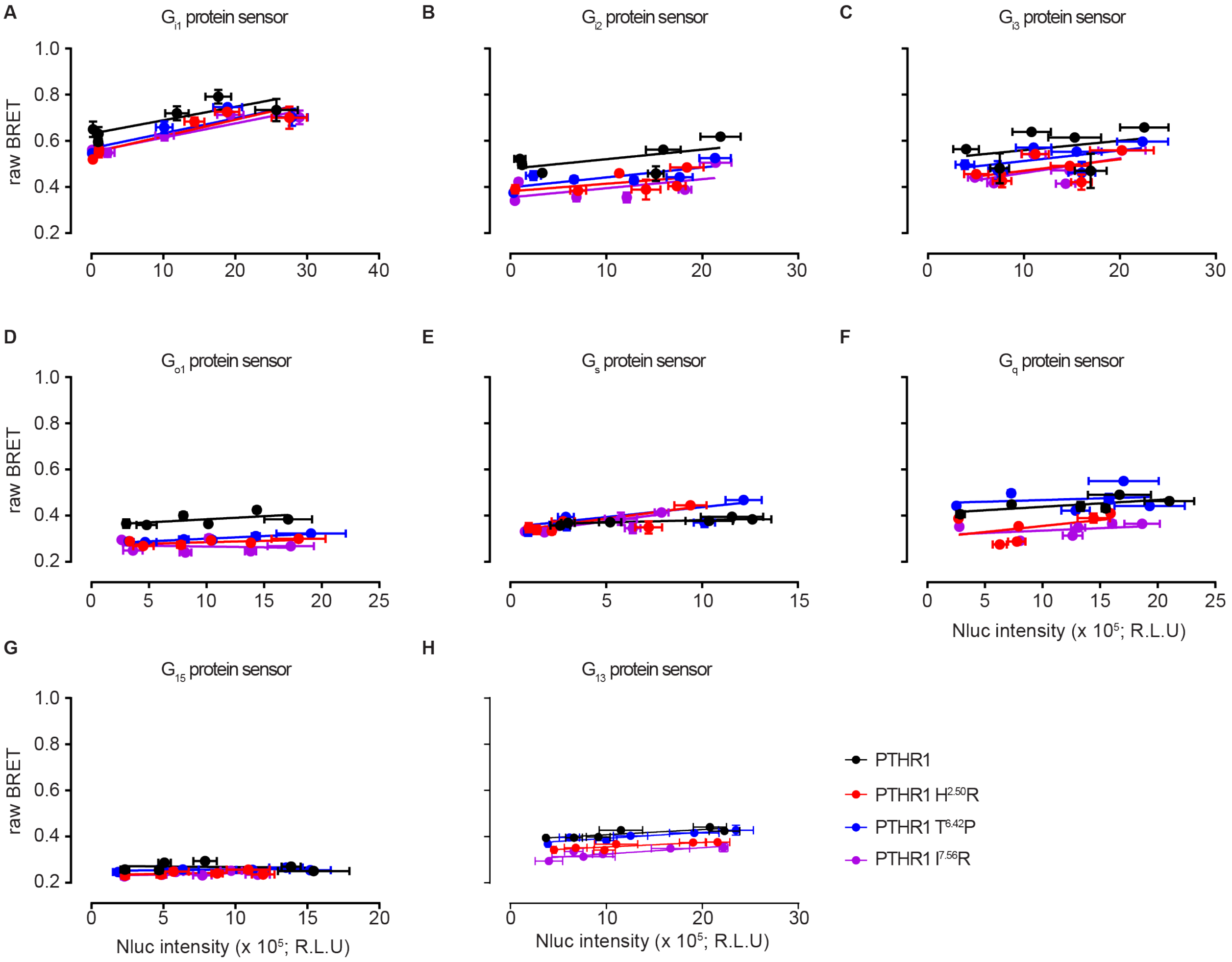


**Fig. S9: BRET over Nluc plots for the assessment of constitutive G protein activity profiles orphan Jansen's disease-related PTHR1 mutants.** BRET over Nluc luminescence plots of HEK293A cells co-transfected with a tricistronic G protein sensor along with wildtype PTHR1 or the indicated point mutants. Data show mean ± s.d. of three individual experiments (two datapoints per experiments at different time points) fitted to a linear correlation.

**Supplementary Table 1:**

**P-values of runs test for deviation from linear correlation in Figure 5.**

| | $G_{i1}$ | $G_{i2}$ | $G_{i3}$ | $G_{o1}$ | $G_{s(short)}$ | $G_q$ | $G_{15}$ | $G_{13}$ |
|---|---|---|---|---|---|---|---|---|
| N1 | 0,7879 | 0.1970 | 0.4909 | 0.6515 | 0.8535 | 0.8247 | 0.2788 | 0.9293 |
| N2 | 0.5333 | 0.8535 | >0.9999 | >0.9999 | 0.7879 | 0.7879 | 0.5333 | >0.9999 |
| N3 | >0.9999 | 0.9545 | 0.7455 | 0.9870 | 0.7879 | >0.9999 | 0.8535 | 0.1970 |
